# Four-dimensional label-free live cell image segmentation for predicting live birth potential of mouse embryos

**DOI:** 10.1101/2024.09.25.614861

**Authors:** Taichi Kanazawa, Tatsuma Yao, Sora Takeshita, Tatsuki Hirai, Ryo Suenaga, Takahiro G Yamada, Yuta Tokuoka, Kazuo Yamagata, Akira Funahashi

## Abstract

Selection of high-quality embryos is critical in assisted reproductive technology (ART), but it relies on visual assessment by experts, and the birth rate remains low. We previously developed a deep learning method to predict the birth of mouse embryos by quantifying the morphological features of cell nuclei. This method involves cell nuclear segmentation on fluorescence microscopy images, but fluorescence labeling of nuclei is not feasible in medical applications. Here, we developed FL^2^-Net, a nuclear segmentation method for time-series three-dimensional bright-field microscopy images of mouse embryos without fluorescence labeling. FL^2^-Net outperformed existing state-of-the-art segmentation methods. We predicted the birth potential of mouse embryos from the nuclear features quantified by bright-field microscopy image segmentation. Birth prediction accuracy of FL^2^-Net (81.63%) exceeded those of other methods and experts (55.32%). We expect that FL^2^-Net, which can quantify nuclear features of embryos non-invasively and with high accuracy, might be useful in ART.

## 2 Introduction

Embryo selection markedly affects birth rate in assisted reproductive technology (ART), one of the most common treatments for infertility currently affecting 186 million people worldwide [1]. In ART, a single embryo with a high potential for live birth is selected out of several embryos fertilized and cultured in vitro through visual assessment of morphological features by embryologists on the basis of grading criteria [2, 3]. For example, the widely used embryo selection based on the Gardner criteria [3] assigns a grade based on the number of the cells in the inner cell mass and trophectoderm, and on the degree of blastocoel expansion at the blastocyst stage, and high-grade embryos are prioritized for transfer. However, substantial worldwide variations in birth outcomes are reported, and the delivery rate per transplanted embryo in ART is 32.1% [4].

The use of machine learning for embryo evaluation is currently attracting widespread interest, because machine learning enables automatic feature extraction and uniform assessment of embryos. Such methods grade human embryos according to existing criteria without bias by experts [5–9]. However, the current grading criteria are defined empirically and may not be the best for predicting the live birth potential of embryos. For example, Khosravi et al. [5] proposed a deep learning method for embryo grading based on Veeck criteria [2]; it achieved an accuracy of 96.94%, but the prediction accuracy of live birth potential was 51.85% [5].

Earlier, we proposed a deep learning ‒ based framework for predicting live birth potential independently of traditional embryo grading criteria [10, 11]. This framework used QCANet, which can perform nuclear segmentation of time-series three-dimensional (3D) fluorescence microscopy images to quantify multivariate time-series data including cell nuclear number, shape, and position [10]. On the basis of these data, the method named Normalized Multi-View Attention Network (NVAN) [11] automatically learns the morphological features distinct between born and aborted embryos and achieves higher accuracy (83.87%) than those of the other methods and experts in birth prediction for mouse embryos. A major challenge for the application of NVAN to human embryos is the lack of a highly accurate method to quantify the multivariate time-series data of nuclei non-invasively. Introducing fluorescent substances into the nuclei is not acceptable for medical applications from an ethical point of view. A high-performance nuclear segmentation method for bright-field microscopy images, which are widely used in medical practice, would be a major step forward.

Deep learning methods have achieved considerable results in microscopy image segmentation [12–17]; in particular, 3D microscopy image segmentation based on deep learning is important in biomedical image analysis [10, 18–21]. For example, StarDist-3D [18] segments an object by approximating it to a star-convex shape; it is most widely used in microscopy image analysis. EmbedSeg [19] is based on clustering and has achieved state-of-the-art performance in seven 3D open-source datasets. QCANet [10] has achieved state-of-the-art performance in 3D fluorescence microscopy images of mouse embryos; it consists of two subnetworks (one for prediction of segmentation masks and the other one for prediction of nuclear centers), it uses the watershed algorithm to partition the region and to generate an instance segmentation image. However, these methods use fluorescence microscopy images with clean signals, not noisy bright-field microscopy images that give signals not limited to target structures [22]. Accurate nuclear segmentation during embryogenesis is particularly difficult, because the shape, number, and size of nuclei and their densities vary with time. In human embryos, segmentation of differential interference contrast (DIC) or Hoffman modulation contrast (HMC) microscopy images has been attempted for specific structures such as the cytoplasm, zona pellucida, blastocoel cavity, and inner cell mass at the pronuclear stages [23] or the blastocyst stage [24–27], but it is necessary to quantify the morphological features of cells or nuclei across all developmental stages.

In this study, we developed Four-dimensional Label-Free Live cell image segmentation Network (FL^2^-Net), an instance segmentation method for 3D bright-field microscopy images of preimplantation mouse embryos. We extracted the time variation in morphological features (such as the number and volume of nuclei) and predicted the live birth potential of mouse embryos on the basis of the extracted multivariate time-series data by using NVAN [11].

## 3 Results

### 3.1 Evaluation of instance segmentation performance

A flow diagram from time-series 3D bright-field images to FL^2^-Net (this study) and NVAN [11] to predict the live birth potential of mouse embryos is shown in Fig. 1. FL^2^-Net consists of a backbone network and three output branches (Fig. 2). The backbone network is based on 3D U-Net [28] and has an encoder ‒ decoder architecture with skip connections. Each skip connection has a Convolutional Gated Recurrent Unit (ConvGRU) [29] for propagating information from the previous frame in the time-series data. This mechanism enables the backbone network to exploit the spatiotemporal features of preimplantation development. The features extracted by the backbone network are transformed into a segmentation mask image through Segmentation Branch, into an energy map through Distance Transformation Branch, and into a central marker image through Detection Branch. In post-processing, these three model outputs are used by the watershed algorithm. The watershed algorithm is an image processing technique used to separate different object areas of a mask image and is used for accurate partitioning of such regions [10, 30–32]. In this algorithm, the intensity in the energy map is treated as an elevation, and the watershed is found where the water body originating from the marker area to the center intersects with the water body from other marker areas. This watershed divides the masked area.

**Fig. 1.**
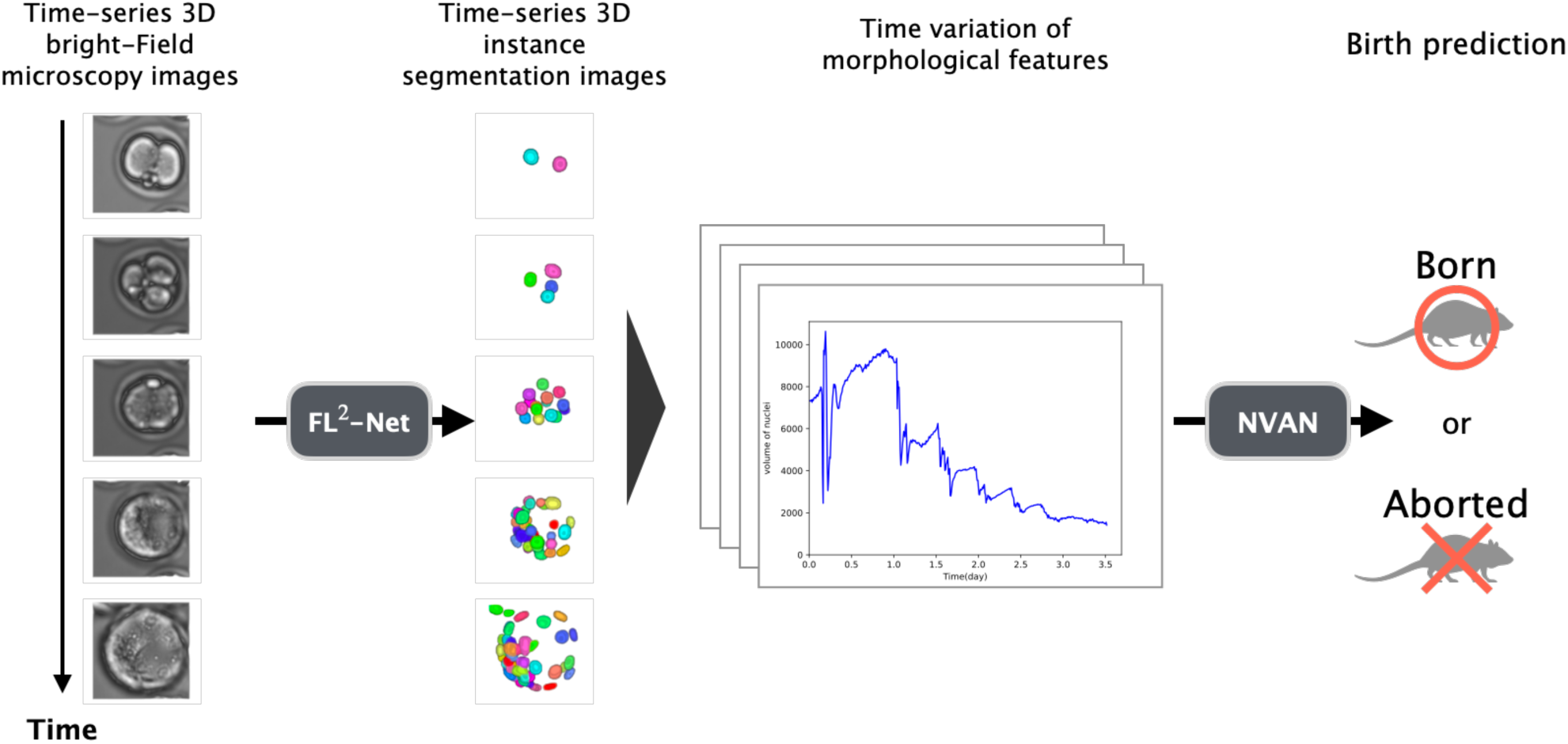
Flow diagram of four-dimensional label-free live cell image segmentation by newly developed FL^2^-Net method to predict live birth potential of mouse embryos. Instance segmentation was performed on each frame of the input time-series 3D bright-field microscopy images. Temporal changes in 11 morphological features, such as the numbers and volumes of nuclei, were extracted, quantified and used as the basis for live birth prediction of mouse embryos with the previously developed NVAN method [11].

**Fig. 2.**
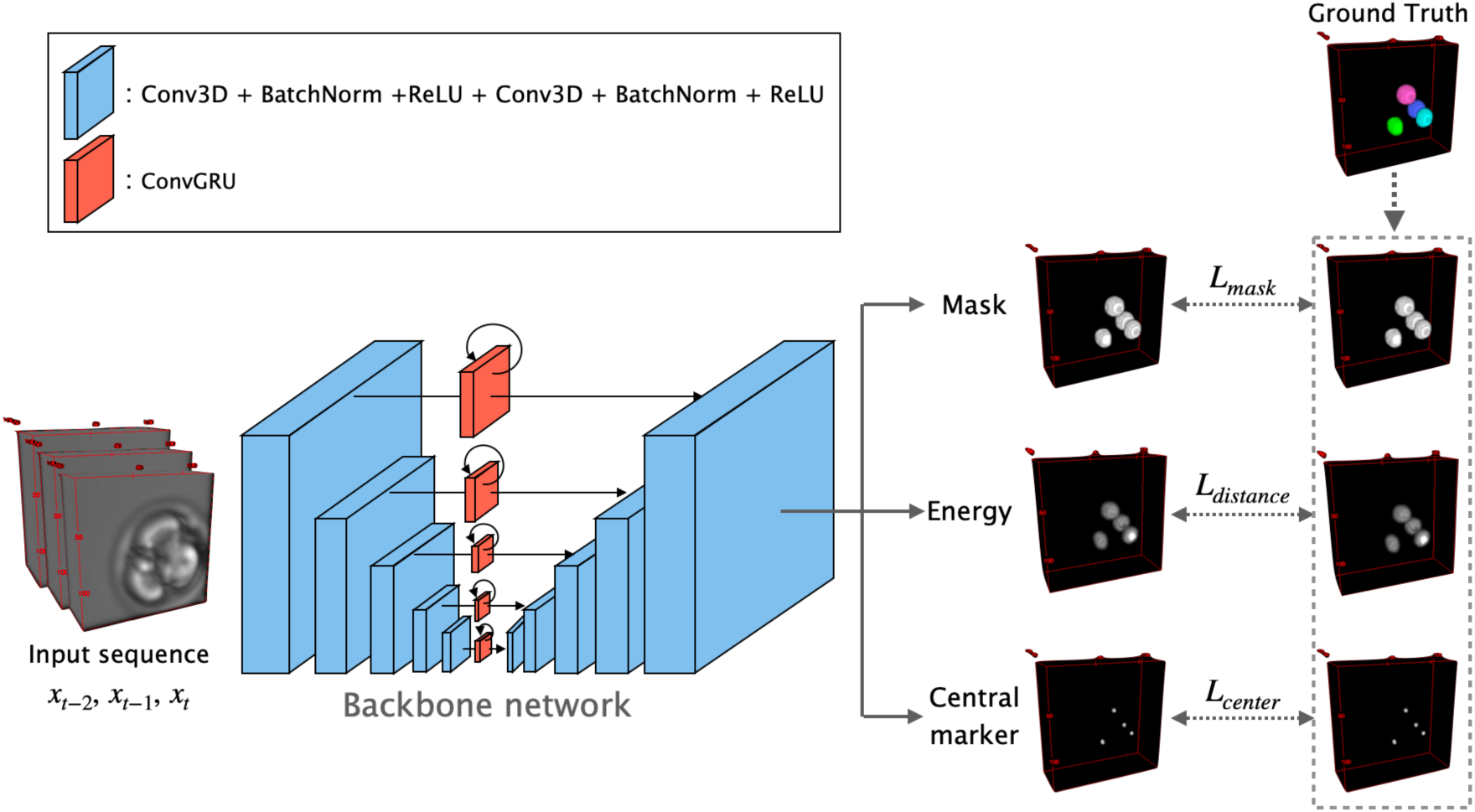
Flow diagram of model training. FL^2^-Net consists of a 3D U-Net-based backbone network and three output branches. The network consists of iterative 3D convolutional layers followed by batch normalization and ReLU activation. Each skip connection in the backbone network has a ConvGRU layer through which the information of previous frames in the sequence is propagated. The model receives an input sequence and outputs a segmentation mask, energy map, and central marker image through the backbone and output branches. In the training phase, the three images are generated from the given ground truth, and each loss (*L*_mask_*, L*_distance_*, L*_center_) is calculated.

To train and evaluate the performance of FL^2^-Net, we created a dataset comprising pairs of time-series 3D bright-field microscopy images and ground truth images. 3D bright-field microscopy images were taken of 84 early mouse embryos from the pronuclear stage to the blastocyst stage every 10 min over day 3.5 after fertilization. A total of 43,504 images were taken. The ground truth images for instance segmentation were created semi-automatically on the basis of fluorescence microscopy images of the same embryos (Supplementary Fig. 1). We randomly divided the entire dataset into training, validation, and test datasets at a ratio of 4:1:1. We compared the segmentation performance of FL^2^-Net with those of StarDist-3D [18], EmbedSeg [19], and QCANet [10], by using a test dataset composed of 7,084 images (14 embryos, 506 timepoints). We evaluated their segmentation performance with four metrics, Intersection of Union (IoU), SEGmentation (SEG) [33], Mean Unweighted Coverage (MUCov) [34], and AP_dsb_ [15]. IoU evaluates the performance of segmentation tasks in general, not just instance segmentation. MUCov and SEG evaluate the absence of false positives and false negatives in instance segmentation. AP_dsb_ detects accuracy at IoU thresholds ranging from 0.1 to 0.9 and evaluates instance segmentation performance considering the trade-off between detection performance and the accuracy of segmentation masks. At a low IoU threshold, AP_dsb_ represents the detection performance when some shape inaccuracy is allowed, whereas at a high IoU threshold it represents the detection performance only when accurate shape prediction is allowed.

FL^2^-Net outperformed all other segmentation methods in IoU, SEG, and MUCov metrics (Table 1). All methods had a large standard deviation of segmentation performance because the accuracy varied greatly depending on the developmental stage. Although IoU decreased over time, FL^2^-Net maintained the highest IoU at all time points (Supplementary Fig. 2). In particular, from day 2.5 (around the blastocyst stage), the IoU of EmbedSeg and StarDist dropped considerably, whereas that of FL^2^-Net remained above 0.5. The highest AP_dsb_ of FL^2^-Net at all IoU thresholds indicated that FL^2^-Net was the most robust method for detecting nuclei (Table 2). Segmentation was relatively accurate for all methods at the early stages of development, but frequent errors appeared at the later stages, when the number of cells increased (Fig. 3). FL^2^-Net accurately captured the nuclear regions at the morula and blastocyst stages, whereas the other methods missed nuclei or incorrectly predicted their shapes (Fig. 4). Overall, FL^2^-Net performed well in terms of both detection and segmentation, and it was the most robust at different developmental stages.

**Fig. 3.**
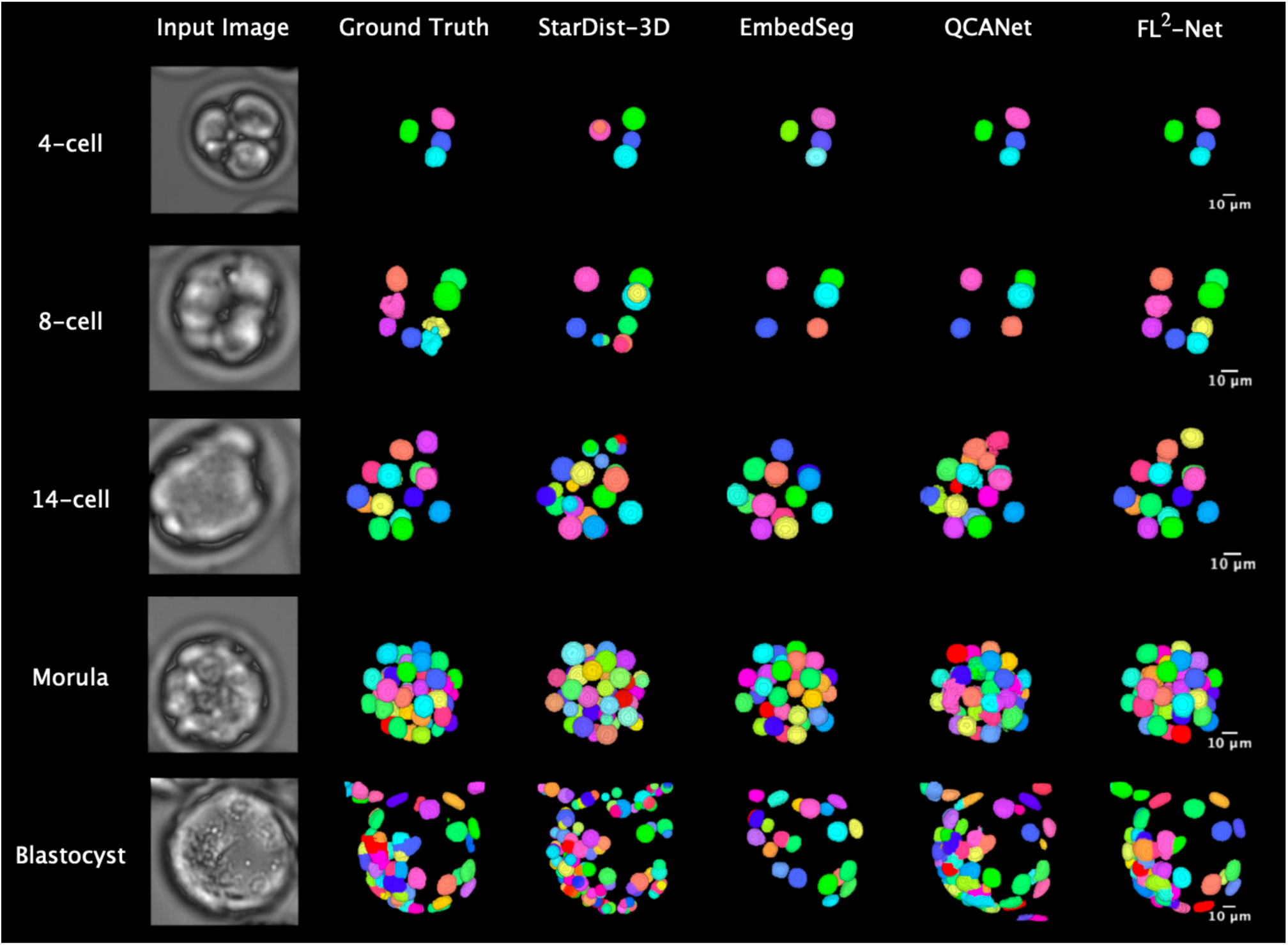
Qualitative comparison of segmentation performances of StarDist-3D [18], EmbedSeg [19], QCANet [10], and FL^2^-Net (this study). Segmentation of 3D bright-field microscopy images of mouse embryos at five developmental stages is shown. Each color represents an individual segmented nuclear region.

**Fig. 4.**
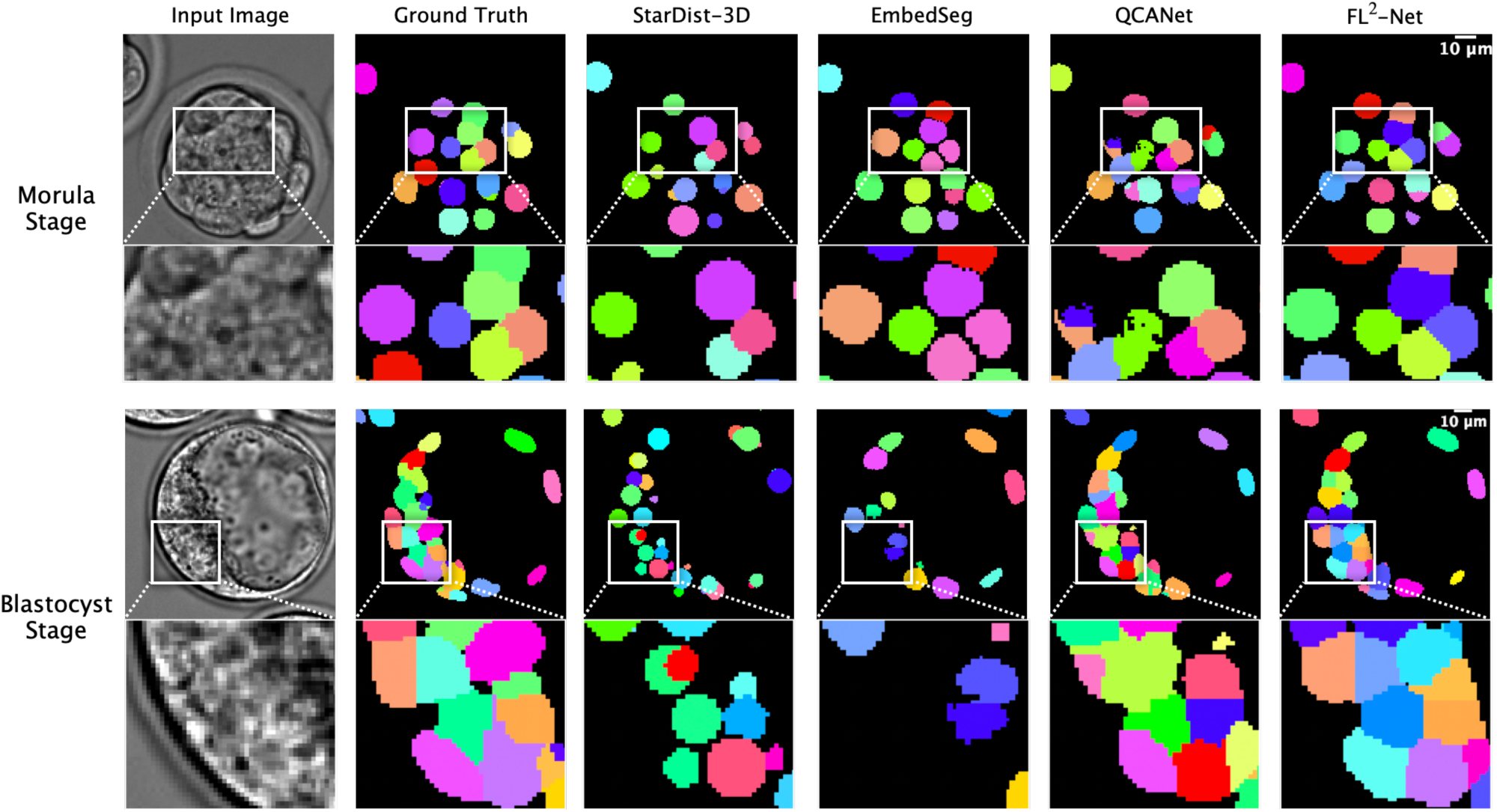
Qualitative comparison of segmentation performances in crowded embryos. Segmentation of 3D bright-field microscopy images of mouse embryos at the morula and blastocyst stages is shown. Each image is a single slice extracted from a 3D image. Each color represents an individual segmented nuclear region.

**Table 1.** Segmentation performances of different methods with the IoU, MUCov, and SEG metrics. Each value (mean and standard deviation) was calculated on the test dataset. Bold, the best score; underline, the second-best score.

|  | IoU | MUCov | SEG |
| --- | --- | --- | --- |
| FL<sup>2</sup>-Net | **0.763 (0.108)** | **0.730 (0.156)** | **0.691 (0.181)** |
| QCANet | <u>0.713</u> (0.128) | 0.674 (0.182) | <u>0.634</u> (0.199) |
| EmbedSeg | 0.682 (0.181) | <u>0.718</u> (0.141) | 0.631 (0.219) |
| StarDist | 0.593 (0.119) | 0.311 (0.210) | 0.413 (0.215) |

**Table 2.** Segmentation performances of different methods with AP_dsb_. Each value was calculated on the test dataset. Bold, the best score; underline, the second-best score.

| IoU threshold | 0.1 | 0.2 | 0.3 | 0.4 | 0.5 | 0.6 | 0.7 | 0.8 | 0.9 |
| --- | --- | --- | --- | --- | --- | --- | --- | --- | --- |
| FL<sup>2</sup>-Net | **0.866** | **0.861** | **0.852** | **0.836** | **0.808** | **0.763** | **0.677** | **0.454** | **0.023** |
| QCANet | <u>0.848</u> | <u>0.834</u> | <u>0.814</u> | 0.786 | 0.745 | 0.683 | 0.576 | 0.320 | 0.005 |
| EmbedSeg | 0.817 | 0.813 | 0.807 | <u>0.797</u> | <u>0.775</u> | <u>0.728</u> | <u>0.614</u> | <u>0.333</u> | <u>0.007</u> |
| StarDist | 0.569 | 0.551 | 0.513 | 0.458 | 0.389 | 0.291 | 0.134 | 0.006 | 0.000 |

The accuracy of bright-field microscopy image segmentation with FL^2^-Net and fluorescence microscopy image segmentation with QCANet [10] up to day 2.5 (as in [11]) is shown in Supplementary Table 1. FL^2^-Net outperformed QCANet in IoU, MUCov, and SEG. In AP_dsb_, FL^2^-Net outperformed QCANet at higher IoU thresholds and underperformed at lower IoU thresholds (Supplementary Table 2). Despite relatively many false detections in the segmentation of bright-field microscopy images, FL^2^-Net accurately predicted the shapes of nuclei.

### 3.2 Ablation studies of FL^2^-Net

Existing methods use a single frame, whereas FL^2^-Net uses consecutive frames in a time series for segmentation. QCANet [10] predicts the segmentation mask and central marker image for the watershed algorithm, whereas FL^2^-Net also predicts the distance image as an energy map, i.e. all the information needed for this algorithm (Supplementary Fig. 3a). To evaluate the incremental benefit of ConvGRU and Distance Transformation Branch, we constructed and trained the following models:

1. FL^2^-Net without ConvGRUs, which performs segmentation using only a single frame (Supplementary Fig. 3b).
2. FL^2^-Net without Distance Transformation Branch, which performs the watershed algorithm on the basis of the distance-transformed image generated from the predicted mask image (Supplementary Fig. 3c).
3. FL^2^-Net without both ConvGRU and Distance Transformation Branch (Supplementary Fig. 3d).

In IoU, MUCov, and SEG, ConvGRU considerably improved the performance of FL^2^-Net (Table 3). In AP_dsb_, accuracy was unaffected at low IoU thresholds because of the use of time-series information, but it was greatly improved at high IoU thresholds (Table 4). The use of distance map prediction had no considerable effect in IoU and MUCov, but it greatly improved FL^2^-Net performance in SEG, i.e. it reduced false negatives. In AP_dsb_ at all IoU thresholds, the use of distance image prediction considerably improved accuracy. In other words, distance image prediction not only improved shape estimation by achieving accurate region partitioning but also improved the detection accuracy of nuclei.

**Table 3.** Ablation analysis of different components in FL^2^-Net with the IoU, MUCov, and SEG metrics. We compared FL^2^-Net, FL^2^-Net without GRU mechanisms (w/o GRU), FL^2^-Net without Distance Transformation Branch (w/o dist), and FL^2^-Net without both ConvGRU and Distance Transformation Branch (w/o both). Each value (mean and standard deviation) was calculated on the test dataset. Bold, the best score; underline, the second-best score.

|  | IoU | MUCov | SEG |
| --- | --- | --- | --- |
| FL<sup>2</sup>-Net | **0.763 (0.108)** | <u>0.730</u> (0.156) | **0.691 (0.181)** |
| FL<sup>2</sup>-Net w/o GRU | 0.748 (0.113) | 0.726 (0.141) | 0.675 (0.185) |
| FL<sup>2</sup>-Net w/o dist | <u>0.758</u> (0.115) | **0.734 (0.146)** | <u>0.679</u> (0.195) |
| FL<sup>2</sup>-Net w/o both | 0.746 (0.113) | 0.715 (0.156) | 0.665 (0.195) |

**Table 4.** Ablation analysis of different components in FL^2^-Net with AP_dsb_. We compared FL^2^-Net, FL^2^-Net without GRU mechanisms (w/o GRU), FL^2^-Net without Distance Transformation Branch (w/o dist), and FL^2^-Net without both ConvGRU and Distance Transformation Branch (w/o both). Each value was calculated on the test dataset. Bold, the best score; underline, the second-best score.

| IoU threshold | 0.1 | 0.2 | 0.3 | 0.4 | 0.5 | 0.6 | 0.7 | 0.8 | 0.9 |
| --- | --- | --- | --- | --- | --- | --- | --- | --- | --- |
| FL<sup>2</sup>-Net | <u>0.866</u> | <u>0.861</u> | **0.852** | **0.836** | **0.808** | **0.763** | **0.677** | **0.454** | **0.023** |
| FL<sup>2</sup>-Net w/o GRU | **0.870** | **0.863** | <u>0.850</u> | <u>0.829</u> | <u>0.795</u> | 0.738 | 0.642 | 0.405 | 0.012 |
| FL<sup>2</sup>-Net w/o dist | 0.857 | 0.851 | 0.841 | 0.823 | 0.793 | <u>0.747</u> | <u>0.658</u> | <u>0.449</u> | <u>0.019</u> |
| FL<sup>2</sup>-Net w/o both | 0.851 | 0.842 | 0.830 | 0.808 | 0.776 | 0.726 | 0.629 | 0.409 | 0.011 |

### 3.3 Extraction of nuclear features in preimplantation embryo

We applied FL^2^-Net to time-series 3D bright-field microscopy images and quantified the temporal variation of 11 morphological features. Comparison of the number of nuclei and mean nuclear volume (Supplementary Fig. 4) quantified by FL^2^-Net and those quantified from ground truth for a single embryo in the test dataset is shown in Supplementary Fig. 5. In the quantification of nuclear number, the error increased in the late stage of development, but the temporal changes were the same as in the ground truth up to about day 2.5. The average nuclear volume also showed that FL^2^-Net captured its stepwise decrease through cell division as ground truth did.

To quantify features to predict the live birth potential of early mouse embryos, we created a dataset for birth prediction consisting of time-series 3D bright-field microscopy images of embryos labeled as either born (110 embryos) or aborted (37 embryos). We applied FL^2^-Net to the time-series 3D bright-field microscopy images, extracted the 11 multivariate time-series data, and used these data to train NVAN [11]. As in [11], we excluded the data after day 2.5, when segmentation accuracy declined. Multivariate time-series data quantified by StarDist-3D, EmbedSeg, and QCANet were also input to NVAN training. To evaluate the classification accuracy, we randomly divided the entire dataset into training and test sets at a ratio of 2:1, and we evaluated the prediction accuracy by using the test set. Birth prediction using FL^2^-Net (accuracy 0.8163, F-measure 0.8861, AUROC 0.6651, AUPR 0.8591) was most accurate in all metrics (Table 5)̶that is, FL^2^-Net was the best method of extracting morphological parameters for birth prediction from bright-field microscopy.

**Table 5.**
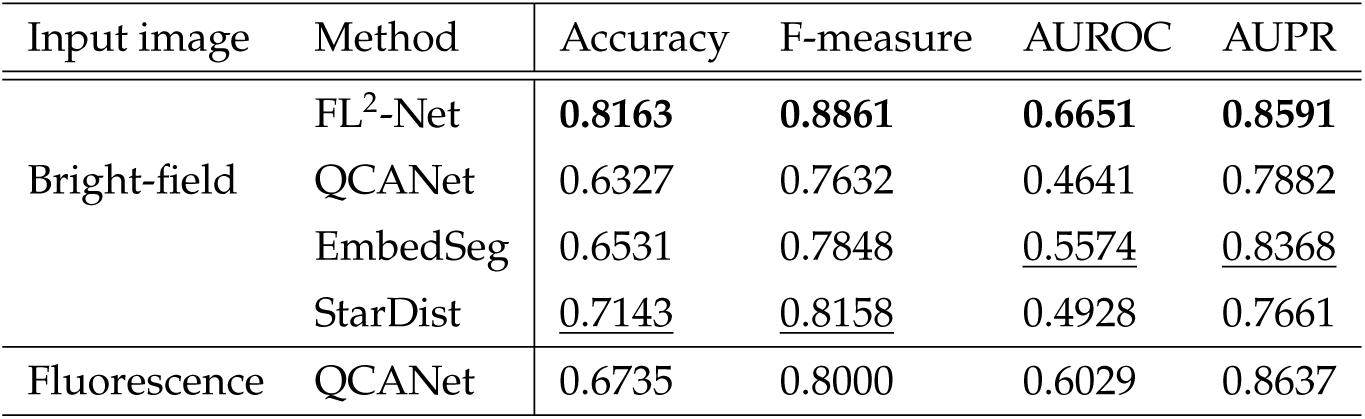
Birth prediction accuracy of each segmentation method. Each value (mean and standard deviation) was calculated on the test dataset. Bold, the best score; underline, the second-best score.

To determine whether it is possible to accurately quantify the morphological features without relying on fluorescence images, we created a dataset for birth prediction consisting of time-series 3D fluorescence microscopy images of the same embryos as the bright-field microscopy images, and compared the accuracy of birth prediction using bright-field microscopy image segmentation with FL^2^-Net and fluorescence microscopy image segmentation with QCANet [10]. FL^2^-Net achieved higher performance for all three indicators except AUPR (Table 5); therefore, high accuracy in birth prediction can be achieved without relying on fluorescence labeling.

### 3.4 Analysis of embryo behavior to predict live birth potential

NVAN with its built-in attention mechanism can visualize the input data variables, which contribute highly to prediction when classifying embryos. In the time-series data of embryos for which live birth potential was correctly predicted, we identified the variables important in prediction by calculating the mean attention value at each time point for each variable. The standard deviation of nuclear volume, the standard deviation of surface area, and the mean nuclear volume were the most important, in that order (Supplementary Fig. 6); the standard deviation of nuclear volume from fertilization to pronucleus fusion contributed highly to birth prediction. The attention map from fluorescence microscopy image segmentation is shown in Supplementary Fig. 7.

In this segmentation, extensive attention was paid to shape information, such as the mean and standard deviation of surface area and standard deviation of aspect ratio; the mean surface area of nuclei after day 1.5 highly contributed to birth prediction. Both the variables and timing focused on in the birth prediction differed greatly between fluorescence microscopy image segmentation and bright-field microscopy image segmentation: the former focused on later stage features, whereas the latter focused on the early stage.

### 3.5 Comparison of prediction accuracy of live birth potential between our method and embryo culture experts

We compared the prediction accuracy of FL^2^-Net with those of 38 embryologists from 13 institutes in Japan (Supplementary Fig. 8). The average accuracy of the experts was 0.5532 *±* 0.0945 (F-measure 0.6680*±*0.1013); it varied widely among the experts, with no correlation with years of experience (Spearman’s *ρ* = *−*0.08, *P* = 0.65), indicating that prediction is difficult even for skilled experts. The accuracy of FL^2^-Net was 0.8163 and the F-measure was 0.8861. Therefore, the ability of FL^2^-Net to extract features for birth prediction far exceeded that of experts.

## 4 Discussion

In this study, we developed FL^2^-Net, an instance segmentation method for time-series 3D bright-field microscopy images of mouse embryos. It outperformed StarDist-3D [18], EmbedSeg [19], and QCANet [10]. FL^2^-Net quantified features of nuclei during preimplantation development, on the basis of which NVAN [11] predicted birth with the highest accuracy among the existing methods. Birth predictions by using the combination of FL^2^-Net and NVAN also outperformed visual assessment by experts in embryo culture; therefore, this combination enables the highly accurate automatic assessment of embryo quality without the need to rely on subjective expert opinion. The main contribution of this study is that FL^2^-Net was validated with brightfield microscopy images taken without fluorescence labeling of the cells. Therefore, FL^2^-Net has potential for medical applications in human embryos.

Structures within the nucleus, such as the nucleolus, which has high molecular density, and the nuclear envelope can also be seen in bright-field images, but the low contrast and high noise make it difficult to identify the nuclear region accurately, especially when the number of embryonic cells increases. StarDist had the lowest performance, probably because the nuclear shapes could not be approximated as star-convex, which is the shape predicted by this method. The significant decrease in AP_dsb_ at high IoU thresholds may have been caused by the inaccuracy of the star-convex approximation (Table 2). The performance of EmbedSeg decreased considerably when the cells became more dense, as in the blastocyst stage; many nuclear regions were missed in areas with high density of the nuclei (Figs. 3,4). EmbedSeg outputs an offset vector for each voxel and implicitly learns that voxels belonging to the same object should point to regions close to each other; these errors are likely caused by ambiguity in the prediction of which object each voxel belongs to at high object densities.

QCANet, like FL^2^-Net, uses the watershed algorithm, but its predicted images had considerable shape inaccuracies. This is partly because QCANet learns segmentation and detection independently in two different models, whereas other methods learn all outputs in a single model. Jointly learning detection and distance map estimation at the same time as segmentation enables more global information such as shape and location to be captured and contributes to the robustness of segmentation prediction [35–37]. The introduction of distance map prediction improved not only the shape estimation but also the detection of nuclear regions (Table 4).

In this study, we attempted segmentation by using spatiotemporal features, whereas other methods̶for instance, segmentation of 3D microscopy images̶process a single frame as input. The use of spatiotemporal features in bright-field microscopy images, which provide few clues for analysis, could improve performance (Tables 3,4). FL^2^-Net with multiple frames referenced by ConvGRU achieved higher performance than when only a single frame was referenced; referring to information in successive frames and making predictions based on consistency in the time-series makes the performance more robust in the case of noisy images.

FL^2^-Net with bright-field microscopy image segmentation predicted birth better than did QCANet with fluorescence microscopy image segmentation (except AUPR, Table 5), but both the variables and the timing of attention in the birth prediction differed greatly between these methods. Interestingly, we found that high attention was paid to nuclear size from fertilization through 2-cell stage when bright-field microscopy image segmentation by FL^2^-Net was used. This finding indicates that some factors may help to predict live birth potential in this period. In ART, embryos are often cultured to the blastocyst stage to identify those with maximum birth potential, because it is believed that only viable embryos can reach this stage [38]. However, the extended culture may have adverse effects on embryos [39], and the choice needs to be made as early as possible. It would be worthwhile to focus on the early stages for birth prediction. In addition, many of the morphological features in this study were of low attention or were not considered at all; therefore, performance could be further improved by excluding them.

In recent years, several studies have proposed deep learning methods to predict pregnancy or birth by using human embryos, and some have shown promising results [40, 41]. However, it is debatable whether these predictions are accurate enough for embryo selection; all of the studies focused on improving prediction accuracy and lacked explainability. Explainable predictive systems are important both in gaining credibility with patients and in supporting experts in their decisions [42]. In this respect, the FL^2^-Net ‒ NVAN combination might be a promising tool for optimal embryo selection in ART.

At present, embryo quality assessment by using FL^2^-Net ‒ NVAN has some limitations for medical applications. (1) The segmentation accuracy may depend on the imaging conditions. Because segmentation by FL^2^-Net is based on four-dimensional features in time-series 3D bright-field microscopy images, its performance may be degraded if the temporal or spatial resolution (especially the number of stacks in the z-axis direction) is low. The image data widely used in medical practice today have lower spatial and temporal resolution than those used in this study. Future work may explore possibilities to increase the robustness against such differences in imaging conditions. Potential future directions include training with images captured at various temporal and spatial resolutions, or the application of super-resolution technology. (2) This study was validated in mouse embryos. Despite similarities in the early embryogenesis of mice and humans, a number of features differ, such as developmental rate and oocyte size [43]. To establish a birth prediction method for human embryos, additional training data are needed for segmentation.

## 5 Methods

### 5.1 Animals

All animal experiments conducted according to the Guide for the Care and Use of Laboratory Animals, and were approved by the Animal Care and Use Committee of the Research Institute of Kindai University (permit number: KABT-2022-012). ICR or B6D2F1 (BDF1) strain mice were obtained from Japan SLC, Inc. ICR males were 15 ‒ 19 weeks old and females were 12 ‒ 16 weeks old; BDF1 males were 11 ‒ 12 weeks old and females were 12 weeks old. Room conditions were standardized (23*^◦^*C, relative humidity 50%, 12-h/12-h light/dark cycle), and mice had free access to water and commercial food pellets. Mice used for experiments were sacrificed by cervical dislocation.

### 5.2 Dataset for segmentation model training and evaluation

The dataset for segmentation model training and evaluation had 43,504 time-series images of 84 early mouse embryos; the images were taken by bright-field microscopy over a period of approximately 3.5 days from immediately after fertilization (pronuclear stage) to the blastocyst stage. Because of difficulties in creating the ground truth of instance segmentation for model training and evaluation from bright-field microscopy images, we used fluorescence microscopy images of the same embryos (Supplementary Fig. 1) injected with histone H2B-mCherry mRNA at the pronuclear stage [44]. The imaging conditions are shown in Supplementary Table 3. To create ground truth, we used QCANet [10], which had been trained to segment nuclei from fluorescence microscopy images. We manually corrected the segmentation errors in the output images of QCANet, mainly in the early stages, by using the image processing platform Fĳi [45]. For 10,172 volumes with significant errors (mainly at the blastocyst stage), Global Walkers, Inc. manually annotated nuclear centroids by using napari [46]. On the basis of the detected nuclear centroids and segmentation images of QCANet, ground truth images were created by using the watershed algorithm. The quality of the corrected images was ensured by checking the correspondence of the nuclei in the pre/post-frames in the time-series images.

### 5.3 Dataset for birth prediction model training and evaluation

Time-series fluorescence and bright-field microscopy images of mouse embryos labeled as born or aborted were taken during the implementation of a previously reported single-embryo transfer experimental system [47]. The imaging conditions were the same as for the segmentation dataset (Supplementary Table 3). On day 4 after in vitro fertilization of oocytes, a single blastocyst from an ICR mouse and six blastocysts from BDF1 mice (as carrier embryos) were transferred into a pseudopregnant ICR female. On day 18.5 after fertilization, the presence of a pup was assessed by cesarean section. ICR and BDF1 mice have different eye colors, so their progenies are easily distinguishable. If no mice were born from carrier embryos, the data were excluded from the analysis, as this indicated problems other than embryo quality. We generated a single embryo transfer dataset consisting of time-series 3D bright-field microscopy images of embryos labeled as either born (110 embryos) or aborted (37 embryos).

### 5.4 FL^2^-Net: proposed segmentation method

The goal of instance segmentation is to identify and segment pixels that belong to each object instance. Instance segmentation is particularly difficult for densely packed objects̶for example, cells at the blastocyst stage. FL^2^-Net is composed of a single backbone network and three output branches (Fig. 2); the backbone refers to previous frames in the time-series for predictions with spatiotemporal features. The three branches output a segmentation mask, energy map, and central marker image. In post-processing, the instance segmentation image is obtained by region partitioning using the watershed algorithm.

#### Backbone network

FL^2^-Net has a backbone network based on 3D U-Net [28], which is used mainly for semantic segmentation tasks. In this network with an encoder ‒ decoder architecture, the image is restored by the decoder on the basis of its features abstracted by the encoder. Multiple skip connections between the encoder and the decoder allow multi-scale feature maps extracted by the encoder to be propagated to the decoder. The backbone network is structured by introducing a ConvGRU [29] into each skip connection of 3D U-Net to propagate spatiotemporal features. GRU [48], a type of recurrent neural network, is a mechanism for selecting and using temporally propagated information; ConvGRU can use spatiotemporal information by replacing the gate operation in the GRU with the convolutional operation.

In FL^2^-Net, to obtain an output frame **y***_t_* from a target frame *x_t_ ∈* R*^D×H×W^* in a time-series image, not only **x***_t_* but also previous frames **x***_t−T_, …,* **x***_t−_*_1_ are used (in this work, *T* = 2). Here *D, H,* and *W* denote the depth, height, and width of the frame, respectively, and **x***_t_* denotes the *t* frame in a time-series image. First, each frame **x***_n_* in an input sequence (**x***_t−T_,* **x***_t−T_* _+1_*, …,* **x***_t_*) is independently input to the same encoder, and a multi-scale feature map **z***_n_* = *{***z***^{l}^|l* = 1, 2*, …L}* (in this work, *L* = 5) is generated. In ConvGRUs, the state is updated by using **z***^{l}^*(*n < t*) extracted from previous frames. When **z***^{l}^_t_* is input to ConvGRU, a feature map **z***^’^_t_* is generated that is based on the information from the previous *T* frames. Similarly to the decoder of 3D U-Net, the multi-scale feature map **z***^’^_t_* is processed stepwise and transformed into a feature map **f***_t_ ∈*R*^C×D×H×W^* with *C* channels. This feature map **f***_t_*is passed to the three output branches.

#### Output branches

All of these output branches consist only of a single 1 *×* 1 *×* 1 convolutional layer followed by a sigmoid function.

#### Segmentation Branch

This branch predicts the probability [0, 1] of belonging to the nuclear region. At inference phase, the output image of the Segmentation Branch is binarized by using a threshold value of 0.5 and used as a segmentation mask in the watershed algorithm. The outputs of the Distance Transformation Branch and Detection Branch are used to partition this mask.

#### Distance Transformation Branch

Each voxel in the distance image predicted by this branch takes the object-specific normalized value of the shortest distance to the background region if it belongs to the nuclear region, or 0 if it belongs to the background region. At inference time, the inverted sign of the output image of this branch is used as the energy map for the watershed algorithm.

#### Detection Branch

This branch predicts the marker image, indicating only the central regions of the nuclei. The ground truth in the training phase is a heatmap with the central coordinates of the nuclei (*p*^*_x_, p*^*_y_, p*^*_z_*) as its peaks. Heatmap *G ∈* R*^D×H×W^* is generated by a Gaussian kernel [49]:

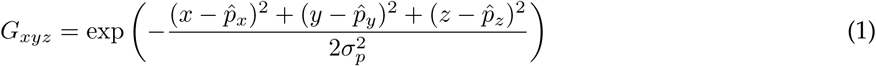

where variance *σ*^2^_p_ is determined as *σ*^2^ = (Size*_x_ ×* Size*_y_ ×* Size*_z_*)*/S* by using the hyperparameter *S* (with *S* = 6) based on object size. Size*_x_,* Size*_y_,* Size*_z_* denote the dimensions of the object in the corresponding directions of (*x, y, z*). At inference time, the output image of the Detection Branch is binarized and labeled and serves as a marker for partitioning the segmentation mask.

#### Model training

The three outputs are trained jointly as a multi-task model, and the loss function for the model (*L*) is calculated as the linear sum of the loss functions for the segmentation (*L*_mask_), distance (*L*_distance_), detection (*L*_center_) images. Dice Loss [50], expressed in the following equation, is used as the loss function for training and is expressed as:

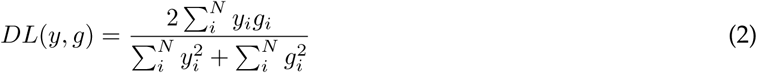

where *y* and *g* are the predicted image and ground truth, respectively, *N* is the number of voxels, and *i* is an index attached to the voxel (*i* = 1, 2*, …, N*). Dice Loss, which evaluates the degree of overlap between prediction and ground truth and is effective when the numbers of voxels in the foreground and background are unbalanced, as in 3D images [50]. Dice Loss was used to ensure stable learning progress despite a large variation in the proportion of nuclei in the embryo images. As an optimizer for model training, we used AMSGrad [51] with a batch size of 2, a training epoch of 10 (141,680 iterations), and an input patch size of 128 × 128 × 128 voxels. To evaluate the segmentation performance, we randomly divided the entire dataset into training and test sets at a ratio of 5:1. The training data were further randomly divided into training and validation sets at a ratio of 4:1.

#### Performance comparison

We compared the performance of FL^2^-Net with those of StarDist-3D [18], EmbedSeg [19], and QCANet [10]. For all previous methods, the hyperparameters of the models had the values recommended in the corresponding papers. For comparison fairness, each model was trained under the same conditions as for FL^2^-Net: AMSGrad optimizer, 10 training epochs, and a 128 *×* 128 *×* 128 voxel patch. Because the GPU memory occupied by different models differs, different batch sizes (4 for StarDist-3D, 8 for EmbedSeg, and 8 for QCANet) were used in view of the GPU memory capacity limit.

#### Evaluation metrics for segmentation performance

We evaluated the segmentation performance with IoU, MUCov [34], SEG [33], and AP_dsb_ [15]. IoU is the ratio of the product set to the sum of the predicted object regions and ground truth regions; it is used to comprehensively measure over- or under-segmentation. MUCov and SEG are metrics for assessing false positives and false negatives in instance segmentation, respectively.

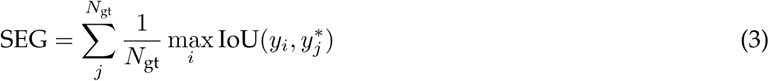

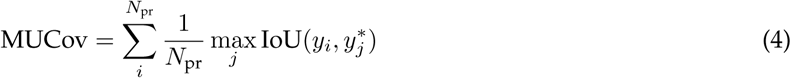

where *N*_gt_ and *N*_pr_ are the number of objects in the ground truth and prediction, respectively, and *y, y^∗^* indicates the set of objects in the ground truth and prediction.

AP_dsb_ at the IoU threshold *τ* means the accuracy in object detection by considering as true positive only those objects whose IoU in the ground truth and predicted images is greater than *τ*. If the objective is to capture the number of objects and their spatial arrangement, a low threshold is sufficient, but to also predict the shape of each object, a high threshold should be used. We used AP_dsb_ at 0.1 increments of *τ* from 0.1 to 0.9, as it is important to comprehensively and accurately quantify embryo morphological features such as the number, volume, and surface area of nuclei.

### 5.5 Birth prediction

We applied instance segmentation to time-series 3D bright-field microscopy images to quantify the time variation in the following morphological features (as in [10]): (1) number of cell nuclei (number), (2) mean volume of a cell nucleus (volume mean), (3) standard deviation of cell nuclear volume (volume sd), (4) mean surface area of a cell nucleus (surface mean), (5) standard deviation of cell nuclear surface area (surface sd), (6) mean aspect ratio of a cell nucleus (aspect ratio mean), (7) standard deviation of cell nuclear aspect ratio (aspect ratio sd), (8) mean solidity of a cell nucleus (solidity mean), (9) standard deviation of cell nuclear solidity (solidity sd), (10) mean distance between the embryo center and the nucleus of every cell (centroid mean), and (11) standard deviation of the distance between the embryo center and the nucleus of every cell (centroid sd). As the quality of the morphological features quantified by different segmentation methods differs, the effectiveness of FL^2^-Net ‒ NVAN combination in birth prediction was assessed by evaluating its performance against birth predictions based on the morphological features quantified by using existing segmentation methods.

#### Birth prediction algorithm architectures and training methods

NVAN [11] is a deep learning method that uses multivariate time-series data quantified by segmentation to predict the live birth potential of embryos without relying on the existing grading criteria. NVAN consists of four components: (1) View-wise Recurrent Encoder, (2) Normalized Hybrid Focus, (3) Multi-view Attention, and (4) View-wise Attentional Feature Fusion. In (1), features for each variable are extracted from time-series multivariate data. In (2), an attention matrix is computed from the extracted features for each variable, in which the relationships between the variants and times are embedded. In (3), the features obtained in (1) are weighted according to the attention matrix. In (4), binary classification is performed on the basis of the weighted features through convolutional layers and fully connected layers. As NVAN aims at binary classification (born or aborted), the loss function in the training was the binary cross entropy loss. The hyperparameters were determined by using Optuna [52], which performed Bayesian optimization. Different multivariate time-series data were quantified for each segmentation method, and the hyperparameters were optimized for each of them.

#### Evaluation for birth prediction performance

The dataset was randomly split at a 2:1 ratio and used as training and test data, respectively. In the training phase, a four-fold cross-validation was performed, and the trained model with the highest F-measure in the validation data was adopted and used to evaluate the test data. The classification performance was evaluated by using the accuracy, F-measure, AUROC (area under the receiver operating characteristic), and AUPR (area under the precision ‒ recall).

#### Classification of embryos by embryo culture experts

Embryologists (38 from a total of 13 institutes) with 1 to 35 years of experience in mouse embryo manipulation classified the early mouse embryos into born/aborted by observing time-series bright-field microscopy images of the single blastocyst transfer dataset (test dataset).

## Data availability

Some of the data used in this paper are available at https://github.com/funalab/FL2-Net. The rest of the data can be provided upon request.

## Code availability

The code used in this paper is available at https://github.com/funalab/FL2-Net.

## Acknowledgements

The research was funded by JSPS KAKENHI, Japan Grant Number 20H03244 to A.F. and JST CREST, Japan Grant Number JPMJCR2331 to A.F. We are grateful to the embryo culture experts at Kindai University, University of Yamanashi, Osaka University, Fukushima Medical University, the University of Tokyo, Kyoto University, Jichi Medical University, Nara Institute of Science and Technology, Fuso Pharmaceutical Industries, Ltd., RIKEN BioResource Research Center, and Central Institute for Experimental Medicine and Life Science for the visual prediction of live birth embryos.

## Author contributions

T.K., Y.T., and A.F. designed the conceptual idea and the study. T.K. implemented the algorithm. T.K., T.G.Y., Y.T., and A.F. designed the methodology and evaluation. T.Y., S.T., T.H., R.S., and K.Y. provided the datasets of mouse embryos. T.K., Y.T., K.Y., and A.F. wrote the manuscript, with suggestions from the other authors.

