## Supplementary materials for "Four-dimensional label-free live cell image segmentation for predicting live birth potential of mouse embryos"

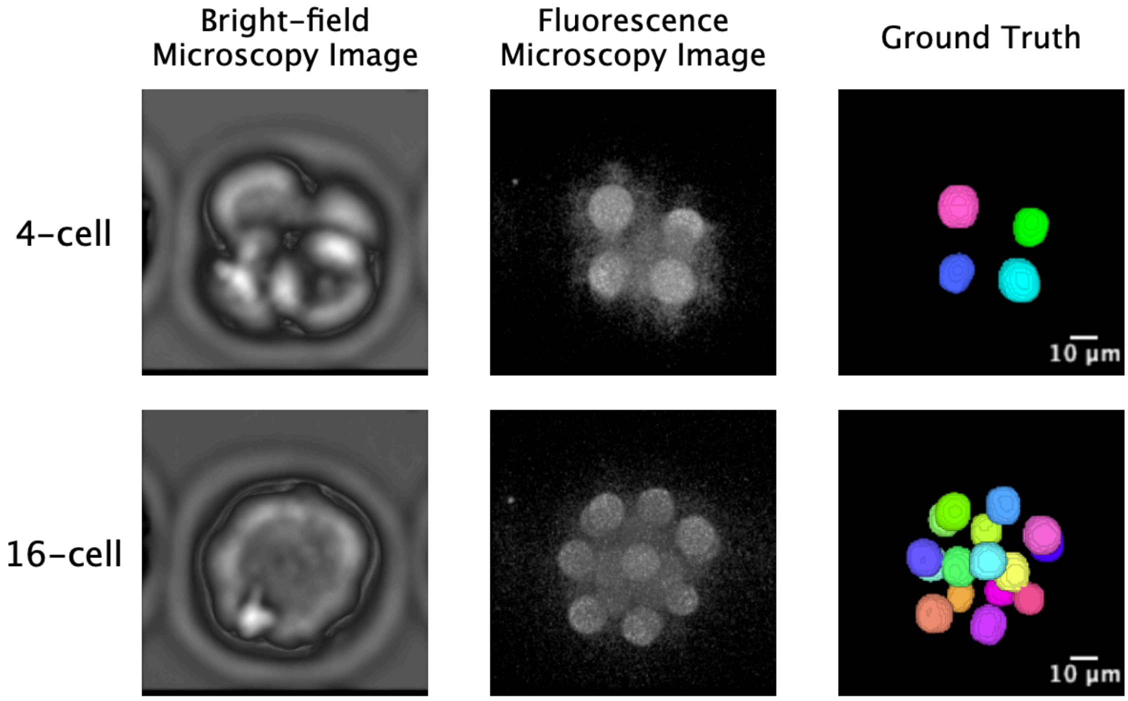


Supplementary Fig. 1 Bright-field and fluorescence microscopy images of the same embryo and the corresponding ground truth.

The ground truth is necessary for learning and accuracy evaluation of the segmentation model; it was created on the basis of fluorescence microscopy images of the same embryo as bright-field images, which were used as input to the model.

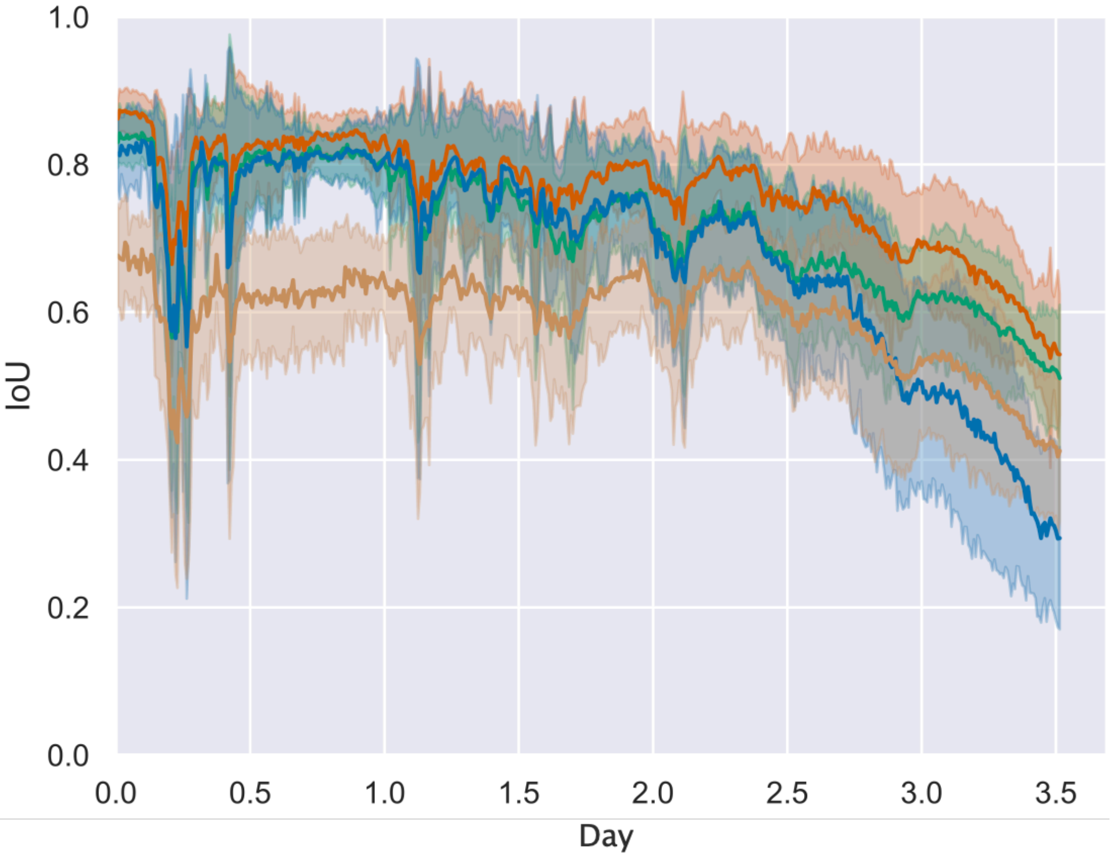


Supplementary Fig. 2 Changes in instance segmentation performance (IoU) with time.

The StarDist-3D [18], EmbedSeg [19], QCANet [10], and FL$^2$-Net methods were used. The horizontal axis corresponds to the time elapsed since fertilization. Each colored line shows the mean value of 14 embryos from the test dataset. Shading represents the standard deviation.

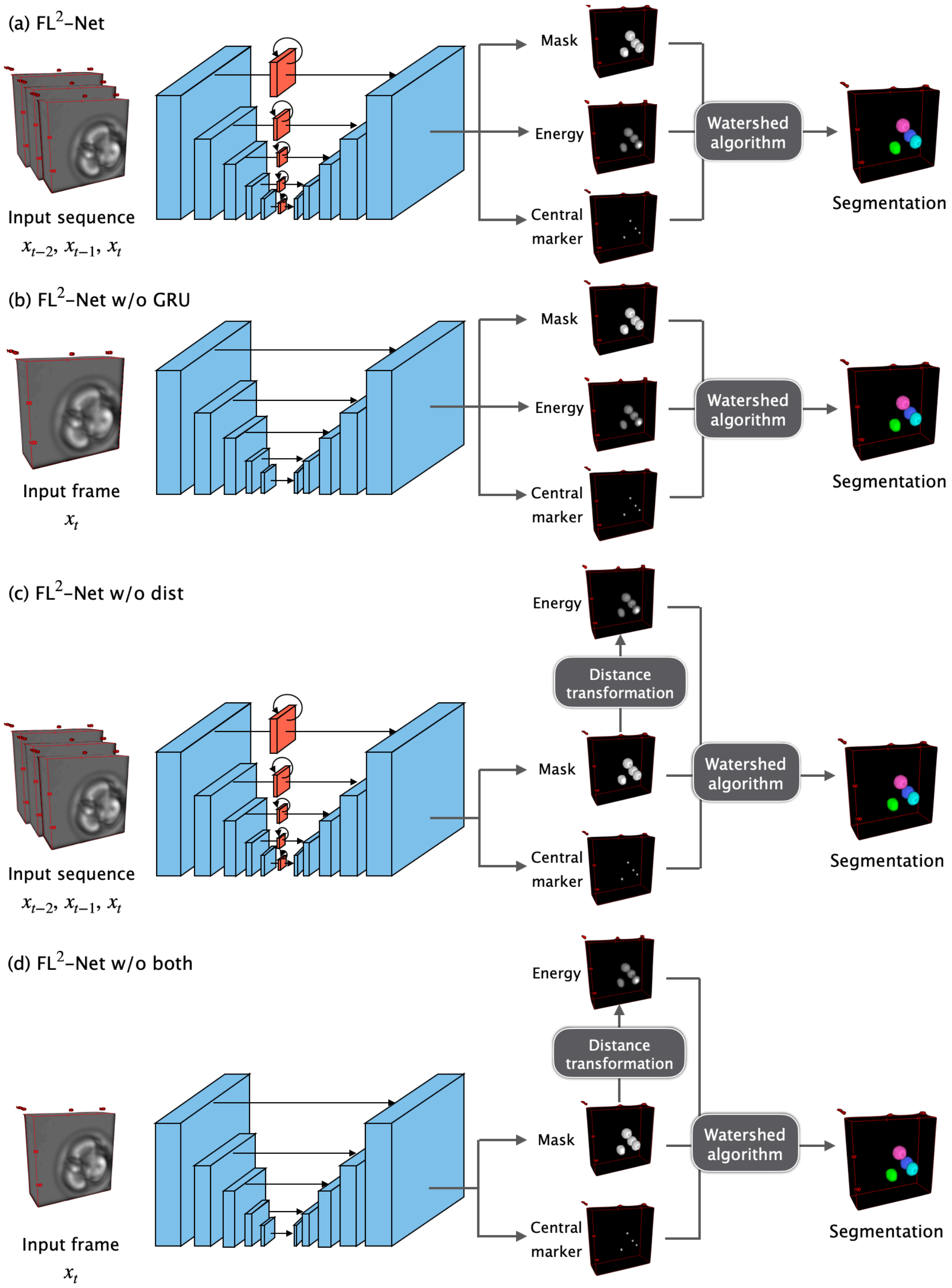


Supplementary Fig. 3 Comparison of inference procedures between FL$^2$-Net and the ablated models.

(a) FL$^2$-Net. (b) FL$^2$-Net without GRU mechanisms (w/o GRU). (c) FL$^2$-Net without Distance Transformation Branch (w/o dist). (d) FL$^2$-Net without both ConvGRU and Distance Transformation Branch (w/o both). In FL$^2$-Net w/o dist and FL$^2$-Net w/o both, as post-processing, distance transformation of the predicted mask image is performed to generate an energy map.

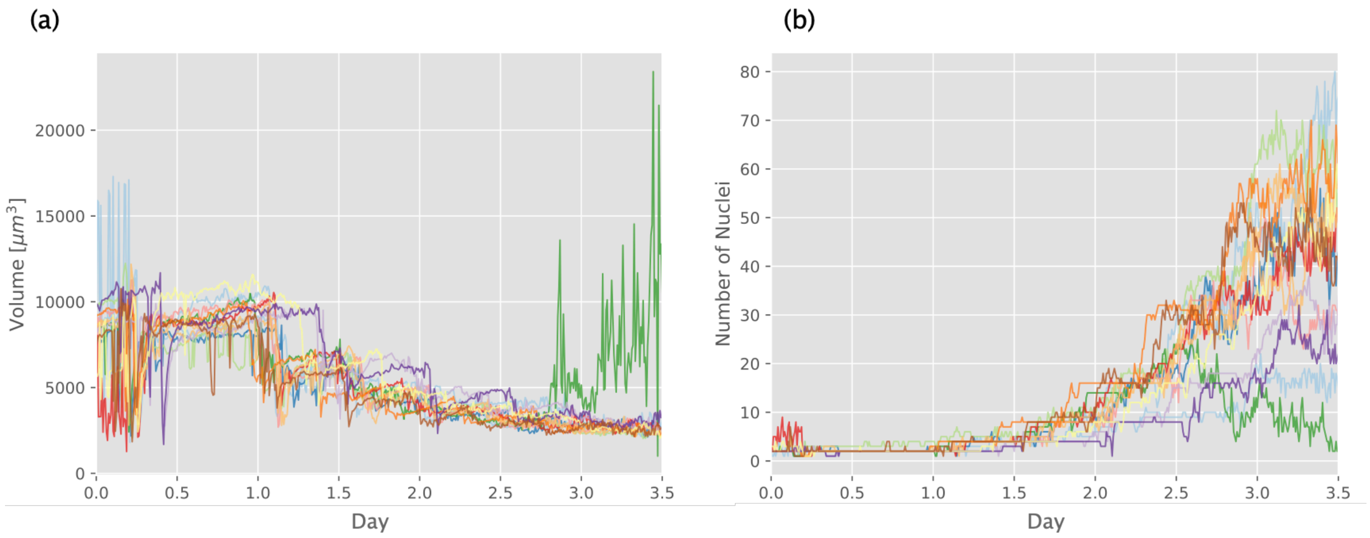


Supplementary Fig. 4 Morphological features of mouse development extracted by FL$^2$-Net from time-series data 3D bright-field microscopy images.

(a) Average volume of a nucleus. (b) Number of nuclei. Each color represents one of the 14 embryos from the test dataset. The horizontal axis represents the time elapsed since fertilization.

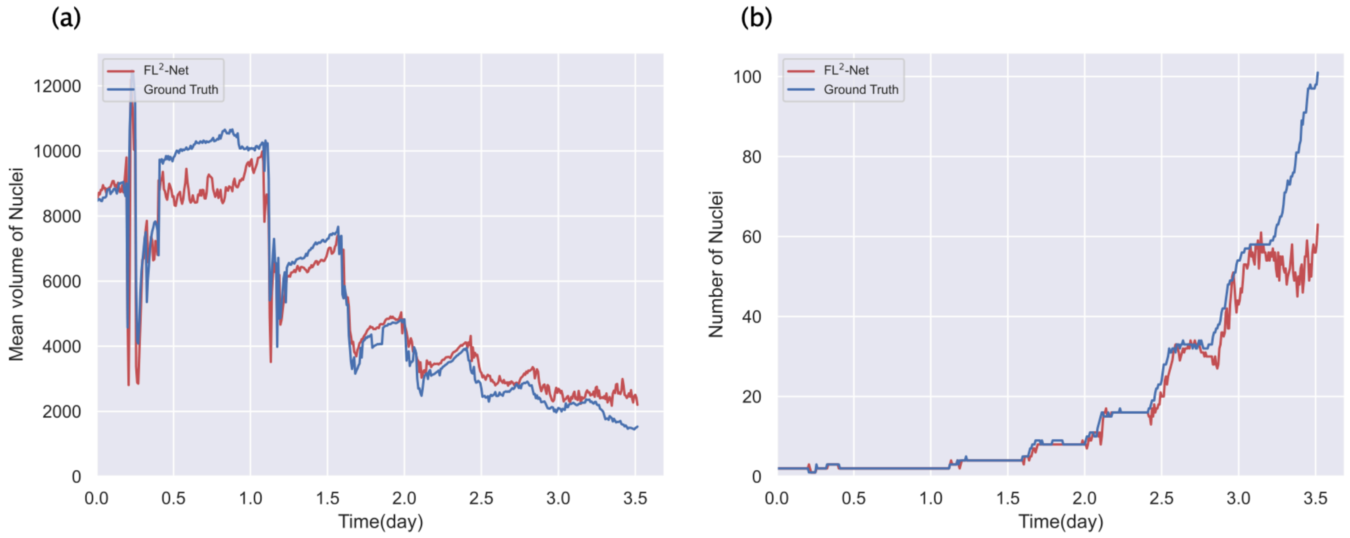


Supplementary Fig. 5 Comparison of morphological features extracted and quantified by $FL^2$-Net with those quantified from ground truth.

Time-series 3D bright-field microscopy images of an embryo from the test dataset were used. (a) Average volume of a nucleus. (b) Number of nuclei. The horizontal axis represents the time elapsed since fertilization.

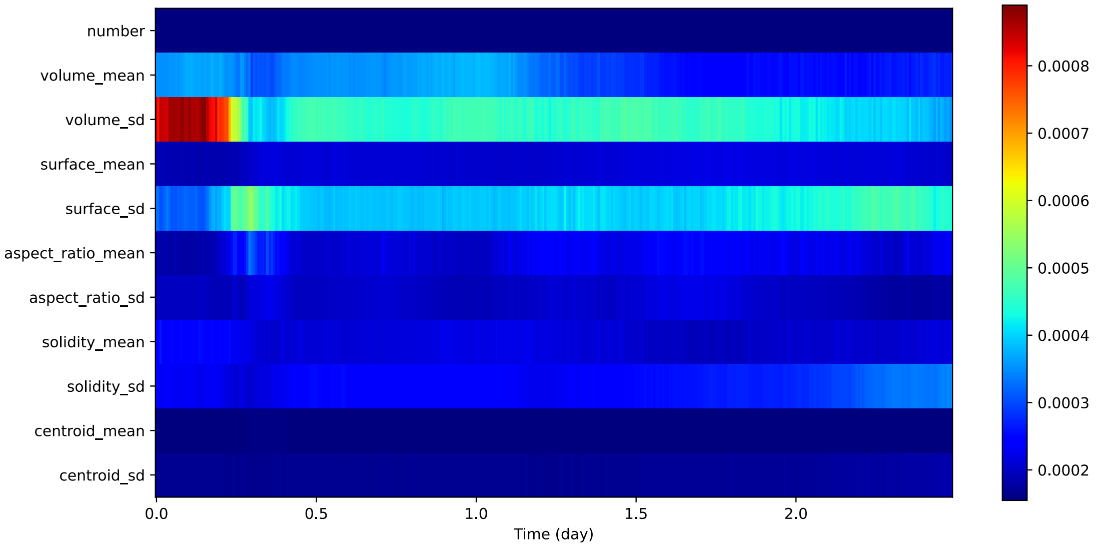


Supplementary Fig. 6 Features quantified by FL$^2$-Net that contributed to the prediction of birth potential.

The figure shows a heat map of the average attention at each time point for each variable in the attention map of the multivariate time-series data quantified by FL$^2$-Net for embryos with correct predictions.

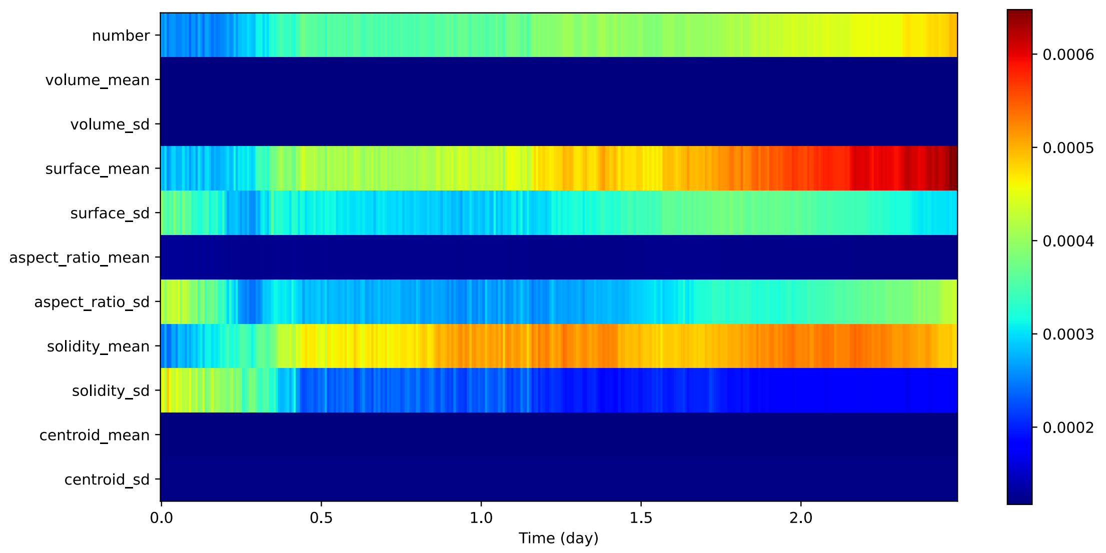


Supplementary Fig. 7 Features quantified by fluorescence microscopy image segmentation with QCANet [10] that contributed to the prediction of birth potential.

The figure shows a heat map of the average attention at each time point for each variable in the attention map of the multivariate time-series data quantified by QCANet [10] for embryos with correct predictions.

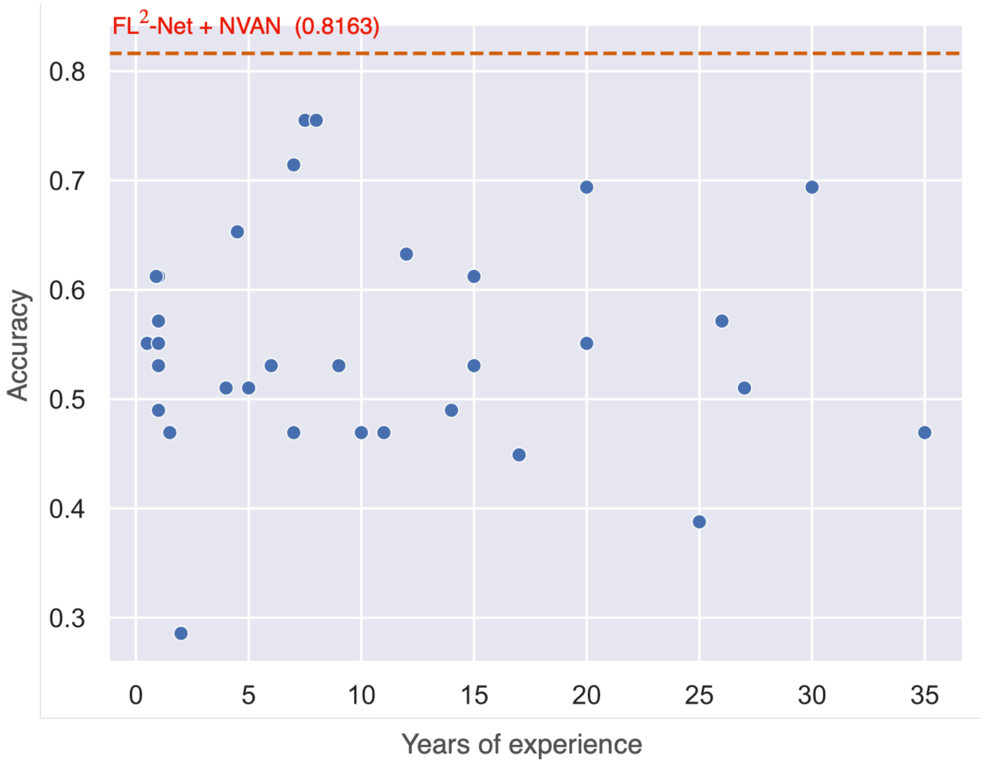


Supplementary Fig. 8    Relationship between number of years of experience in mouse embryo manipulation and prediction accuracy by experts.

The accuracy of birth prediction varied widely among the experts and was not correlated with the number of years of experience (Spearman's $\rho = -0.08$, $P = 0.65$).

Supplementary Table 1 Quantitative performance comparison of bright-field image segmentation (BF segmentation) by FL$^2$-Net and fluorescence image segmentation (FC segmentation) by QCANet [10] using IoU, MUCov, and SEG up to day 2.5. Each value (mean and standard deviation) was calculated on the test dataset. (Bold, the best score)

| | IoU | MUCov | SEG |
|---|---|---|---|
| BF segmentation by FL$^2$-Net | **0.798 (0.086)** | **0.792 (0.107)** | **0.762 (0.128)** |
| FC segmentation by QCANet | 0.768 (0.045) | 0.742 (0.091) | 0.753 (0.057) |

Supplementary Table 2 Quantitative performance comparison of bright-field image segmentation (BF segmentation) by FL$^2$-Net and fluorescence image segmentation (FC segmentation) by QCANet [10] using $AP_{dsb}$ up to day 2.5. Each value was calculated on the test dataset. Bold, the best score.

| IoU threshold | 0.1 | 0.2 | 0.3 | 0.4 | 0.5 | 0.6 | 0.7 | 0.8 | 0.9 |
|---|---|---|---|---|---|---|---|---|---|
| BF segmentation by FL$^2$-Net | 0.928 | 0.927 | 0.925 | 0.921 | 0.909 | 0.882 | 0.813 | **0.578** | **0.030** |
| FC segmentation by QCANet | **0.969** | **0.968** | **0.967** | **0.963** | **0.953** | **0.938** | **0.844** | 0.126 | 0.000 |

Supplementary Table 3 Imaging conditions of mouse embryos for training and evaluation of (a) nuclear segmentation and (b) birth prediction models

| | |
|---|---|
| Observation target | Mouse embryo |
| Microscope and confocal system | CV1000 (Yokogawa Electric Corp.) |
| Laser power [$mW$] | 0.1 |
| Exposure time [$msec$] | 100 |
| Image size [$voxel$] | 512×512×51 |
| Spatial resolution ($x : y : z$) [$\mu m/voxel$] | 0.8 : 0.8 : 2.0 |
| Time resolution [$min$] | 10 |
| Number of time slices | (a) 506 / (b) 509-519 |
| Number of embryos analyzed | (a) 84 / (b) 147 |
